# Lesion-Connectome Mapping of Limb and Oral Apraxia: Distributed vs Focal Networks

**DOI:** 10.64898/2025.12.13.694084

**Authors:** Yoshiyuki Nishio, Ryo Itabashi, Yuka Kataoka, Yukako Yazawa, Etsuro Mori

## Abstract

**Background and Objectives:** To characterize the anatomical network architectures underlying effector-specific apraxias, specifically comparing the neuroanatomical substrates of limb apraxia (LA), buccofacial apraxia (BFA), and apraxia of speech (AOS).

**Methods:** We evaluated 136 patients with acute left-hemisphere ischemic stroke for the presence of LA, BFA, and AOS. We used voxel-based, region-of-interest-based, and connectome-based lesion-symptom mapping to identify the cortical lesions and white matter disconnections associated with each apraxia subtype.

**Results:** LA was associated with damage to the inferior parietal lobule and widespread white matter disconnections, including intrahemispheric temporo-parietal pathways and interhemispheric transcallosal fibers. In contrast, BFA was associated with localized anterior lesions centered on the mid-lower portion of the left precentral gyrus and adjacent frontal operculum, without massive long-range white matter disconnections. While primary analyses for AOS yielded no suprathreshold clusters, likely washed out by the spatial variance of lesions causing concomitant aphasia, exploratory analyses localized AOS to the mid-portion of the precentral gyrus, situated slightly dorsal to the core region of BFA.

**Discussion:** LA and oral apraxias differ fundamentally in their underlying network architectures rather than merely their effectors. LA involves the disruption of a distributed, bilaterally integrated temporo-parietal network. Conversely, oral apraxias depend on localized anterior networks, with a dorso-ventral dissociation within the precentral gyrus distinguishing pure AOS (dorsal primary motor) from BFA (ventral premotor). These findings highlight how the brain uses distinct network principles, distributed versus localized, to control different classes of skilled action.

## INTRODUCTION

Clinical neurology has long recognized apraxias as a distinct class of motor disorders characterized by impairments in skilled action, typically resulting from left-hemisphere damage^1–3^. Clinically, apraxias are categorized by the effectors involved. Limb apraxia (LA) affects upper extremity movements, while oral apraxias are subdivided into buccofacial apraxia (BFA) for non-verbal actions and apraxia of speech (AOS) for motor speech production.

Classical behavioral neurology has established a large-scale anatomical dissociation between these syndromes: oral apraxias are primarily associated with frontal lobe lesions, whereas LA is strongly linked to parietal lobe damage^1,2,4^. However, this classical dichotomy is difficult to reconcile with a simple somatotopic organization of action. If the brain organized skilled movements strictly by effector somatotopy, damage to the frontal motor and premotor cortices should produce both oral and limb apraxias. The fact that frontal lesions typically spare limb praxis, while parietal lesions specifically disrupt it, suggests that these apraxias are governed by distinct principles of network organization rather than simple cortical somatotopy. While previous mass-univariate lesion studies have extensively mapped the cortical correlates of these disorders^5,6^, the contribution of underlying white matter tract disconnection remains insufficiently quantified^7,8^. It remains unclear how structural disconnections within the fronto-parietal system differentially give rise to limb versus oral apraxia.

To address this gap, we conducted a comprehensive lesion-symptom mapping analysis in 136 patients with acute stroke, using voxel-based, region-of-interest-based, and connectome-based approaches. By directly comparing the cortical topographies and structural disconnection patterns of LA and BFA within a single cohort, we aimed to elucidate the distinct network architectures underlying effector-specific apraxias.

## METHODS

### Study population

Participants were retrospectively selected from a cohort of 2,146 consecutive patients with acute ischemic stroke admitted to Kohnan Hospital between April 2007 and March 2012. This cohort is identical to that described in our previous study^9^. Clinical and investigative data were prospectively collected and registered in a standardized database. Diagnosis of ischemic stroke was confirmed by board-certified stroke neurologists based on clinical and neuroimaging findings.

Inclusion criteria were as follows: (1) first-ever stroke onset; (2) hospital admission within 7 days of onset; (3) isolated non-lacunar infarction restricted to the left middle cerebral artery territory, verified by MRI; (4) right-handedness; (5) native Japanese speaker; (6) no history of dementia; and (7) completion of a comprehensive neuropsychological evaluation by speech-language pathologists during hospitalization. Neuropsychological evaluations were indicated for patients exhibiting behavioral or language impairments interfering with daily activities. A total of 136 patients met these criteria (**Table 1**).

**Table 1.** Demographic and clinical characteristics of the study cohort.

|  | **LA**<br>(n = 17) | **BFA**<br>(n = 20) | **AOS**<br>(n = 22) | **No apraxias**<br>(n = 93) | **Total**<br>(n = 136) |
| --- | --- | --- | --- | --- | --- |
| Age (Mean ± SD) | 76.5 ± 11.6 | 69.5 ± 11.0 | 67.0 ± 13.9 | 70.4 ± 12.9 | 70.5 ± 12.9 |
| Sex (Female) | 5 | 2* | 9 | 42 | 55 |
| NIHSS (Mean ± SD) | 13.6 ± 9.6* | 10.9 ± 7.8* | 9.5 ± 8.6* | 5.7 ± 5.8 | 7.3 ± 7.2 |
| Stroke subtype |  |  |  |  |  |
| Cardioembolic | 11 | 7 | 7 | 35 | 53 |
| Large artery atherosclerotic | 4 | 10 | 9 | 34 | 49 |
| Others | 2 | 3 | 6 | 24 | 34 |
| Lesion volume (cm<sup>3</sup>, Mean ± SD) | 86.7 ± 94.9* | 82.5 ± 89.2* | 50.2 ± 89.6 | 19.2 ± 26.1 | 31.2 ± 49.1 |
| Aphasia type |  |  |  |  |  |
| Broca | 5 | 8 | 15 | 0 | 15 |
| Pure AOS | 0 | 1 | 7 | 0 | 7 |
| Wernicke | 9 | 6 | 0 | 19 | 32 |
| Conduction | 0 | 0 | 0 | 1 | 1 |
| Transcortical sensory | 1 | 1 | 0 | 14 | 16 |
| Anomic | 1 | 2 | 0 | 16 | 19 |
| Others | 0 | 2 | 0 | 4 | 6 |
| No aphasia or AOS | 1 | 0 | 0 | 39 | 40 |
| Comorbid apraxia |  |  |  |  |  |
| BFA | 5 | - | 9 | - | 20 |
| AOS | 5 | 9 | - | - | 22 |
| LA | - | 5 | 5 | - | 17 |

Note that apraxia groups are not mutually exclusive; therefore, the sum of patients across subgroups exceeds the total cohort size. \* $p < 0.05$ compared to the 'No apraxia' group.
Abbreviations: AOS, apraxia of speech; BFA, buccofacial apraxia; LA, limb apraxia; NIHSS, National Institutes of Health Stroke Scale.

The study protocol was approved by the Kohnan Hospital Ethics Committee (2015-0107-04). According to the institutional review board policy, individual informed consent was waived due to the retrospective nature of the study. Instead, an opt-out approach was employed, and information regarding the study was publicly disclosed on the hospital website to provide patients with the opportunity to refuse participation. All participant data were anonymized before analysis to ensure patient confidentiality.

### Behavioral assessments

All assessments were conducted by trained speech-language pathologists. The median interval between stroke onset and neuropsychological evaluation was 7 days [IQR: 5–10]. Patients were instructed to execute motor actions in response to verbal commands and imitation. For the purpose of lesion-symptom mapping, the diagnosis of each apraxia subtype was operationalized as a binary variable (present vs. absent).

LA was assessed using symbolic actions of the left upper limb (to avoid confounds from right hemiparesis). Tasks included communicative gestures (military salute, beckoning, waving goodbye) and pantomime of tool use (brushing teeth, combing hair, shaving). LA was diagnosed based on the presence of distinct errors, including incorrect movement direction, abnormal hand configuration, or action substitution.

BFA was assessed through the execution of non-verbal oropharyngeal and facial movements (coughing, clicking the tongue, whistling). A diagnosis was made when patients failed to perform these actions or produced substitution errors on verbal command or imitation.

AOS was diagnosed based on specific speech characteristics observed during free conversation, cartoon description, repetition, and reading-aloud subtests of the Standard Language Test of Aphasia (SLTA)^10^. Diagnostic features included slow speech rate, distorted sound substitutions or additions, articulatory groping, lengthened intersegment durations, or syllable segmentation. Aphasia subtypes were also classified according to the SLTA.

### Imaging procedures

Structural brain imaging was acquired using a 1.5-T scanner (Signa Excite, GE Medical Systems, Milwaukee, WI, USA). Lesions were identified on axial T2-weighted images (TR=3,000 ms; TE=80 ms; matrix=320×256; FOV=22×22 mm; thickness=6 mm; gap=2.0 mm) or FLAIR images (TR=8,002 ms; TE=126 ms; TI=2,000 ms; matrix=256×224; FOV=22×22 mm; thickness=6 mm; gap=2.0 mm). The median intervals between stroke onset and MRI, and between neuropsychological evaluation and MRI, were 9 days [IQR: 7–12] and 3 days [IQR: 1–6], respectively.

Native images and binary lesion masks were normalized to a standard T2-weighted or FLAIR template using a nonlinear transformation with enantiomorphic lesion-cost function masking^11^. The resulting images were resampled to 1-mm isotropic voxels.

To quantify white matter network disruption, we employed connectome-based lesion-symptom mapping (CLSM) based on the "indirect structural disconnection" framework^12^. This method estimates structural disconnection by projecting individual lesion masks onto a normative healthy connectome, avoiding artifacts associated with tracking through pathological tissue. Voxel-wise disconnection severity maps were generated using the Network Modification (NeMo) tool^13^. The reference tractography was derived from healthy individuals using advanced parameters (deterministic algorithms with SIFT2 re-weighting) to capture comprehensive white matter architecture.

### Statistical analyses

We performed voxel-based lesion-symptom mapping (VLSM), region-of-interest (ROI)-based lesion-symptom mapping (ROI-LSM) and connectome-based lesion-symptom mapping (CLSM) using a General Linear Model (GLM) framework in the NiiStat toolbox (https://github.com/neurolabusc/NiiStat)^14^.

#### 1. Analysis of clinical and demographic data

Differences in clinical and demographic characteristics between patients with and without each apraxia subtype were assessed using appropriate non-parametric tests. Furthermore, given the high clinical comorbidity among apraxia subtypes, we performed secondary behavioral analyses on mutually exclusive clinical subgroups (e.g., LA+/BFA- vs. LA-/BFA+ vs. LA+/BFA+) to disentangle their specific clinical profiles, such as stroke severity (NIHSS) and total lesion volume. Importantly, this mutually exclusive subgrouping was used exclusively for descriptive purposes, namely, the clinical profiling in **eTable 1** and the visualization of simple lesion overlaps in **eFigure 1**. All statistical lesion-symptom mapping (LSM) analyses used the inclusive full-cohort design described below.

#### 2. Image modalities and data types

VLSM and ROI-LSM: We used binary lesion masks (lesioned = 1, intact = 0) to assess the impact of focal brain damage at the voxel level.

ROI-LSM: We used continuous measures representing the percentage of lesioned tissue within predefined anatomical regions to assess regional brain damage burden.

CLSM: We used continuous measures of tract disconnection severity (values ranging from 0 to 1) to quantify white matter disruption. Derived from deterministic tractography, this metric represents the proportion of disconnected streamlines relative to the total number of streamlines in a given tract. This continuous variable preserves the graded severity of structural disconnection for regression analysis.

#### 3. Grouping strategy

To investigate the neural correlates of each apraxia subtype, comparisons were made between symptom-present and symptom-absent groups. The symptom-present group was defined inclusively: it comprised all patients exhibiting the specific apraxia subtype, regardless of comorbidity with other apraxia types. For instance, the AOS-present group included patients with AOS without other apraxia subtypes, as well as those with concomitant limb or buccofacial apraxia. This inclusive approach contrasts with analyses restricted to pure cases, allowing us to use the full variability of the cohort in the GLM regression.

#### 4. GLM specifications and significance testing

To identify the unique neuroanatomical correlates of each apraxia subtype while controlling for their high clinical comorbidity, the behavioral variables for LA, BFA, and AOS were entered simultaneously into all GLM analyses. This approach corresponded to a multiple logistic regression for the VLSM, and multiple linear regressions for the ROI-LSM and CLSM. For VLSM, analysis was restricted to voxels damaged in at least 10 patients to ensure statistical power. For ROI-LSM, we calculated the percentage of lesioned tissue within each of the 65 left-hemisphere regions defined by the Johns Hopkins University grey and white matter atlas^15^. We modeled the relationship between lesion location (or disconnection severity) and behavioral deficits (presence/absence of apraxia) while controlling for total lesion volume as a continuous nuisance regressor. Statistical significance was determined using the non-parametric Freedman-Lane permutation method (5,000 permutations)^16^, with a family-wise error (FWE) corrected threshold of *p* < 0.05.

#### 5. Control for hemispheric bias in CLSM

To rigorously control for the non-specific spatial distribution of stroke lesions, we employed a lesion-flipped control approach for the CLSM analysis^17^. The original left-hemisphere lesion masks were flipped across the midsagittal plane to create "virtual" right-hemisphere lesions for all patients. In the GLM analysis, these virtual right-hemisphere cases were all assigned a behavioral score of 0 (symptom absent) for each apraxia subtype. By incorporating this combined dataset into the exact same CLSM pipeline, we forced the model to treat the flipped lesions as asymptomatic controls. This approach allowed us to verify that the identified disconnected networks represent true anatomical associations with the apraxic symptoms, rather than mere artifacts driven by the typical topography of middle cerebral artery strokes.

## RESULTS

### Clinical characteristics

Demographic and clinical characteristics are summarized in **Table 1** and **eTable 1**. Age did not significantly differ between patients with and without apraxias. Male prevalence was significantly higher in the BFA group compared to patients without apraxia (*p* = 0.004). Patients with apraxias presented with greater stroke severity, as indicated by significantly higher National Institutes of Health Stroke Scale (NIHSS) scores across all apraxia subtypes compared to the non-apraxia group (*p* < 0.05 for all comparisons). Regarding lesion volume, patients with LA and BFA had significantly larger lesions than those without apraxia (p = 0.010 and p = 0.005, respectively), whereas patients with AOS did not show a significant difference (p = 0.12). At a descriptive level, language profiles also varied among groups; notably, AOS was exclusively associated with Broca’s aphasia or presented without aphasia.

To better understand the clinical profiles underlying the high comorbidity (over half of patients with one type also exhibited the others), we compared demographic and clinical variables across mutually exclusive subgroups (e.g., LA+/BFA- vs. LA-/BFA+ vs. LA+/BFA+). NIHSS scores did not differ significantly among these mutually exclusive subgroups. When comparing mutually exclusive oral apraxia groups, patients with BFA-/AOS+ had significantly smaller lesion volumes than those with BFA+/AOS- (p = 0.009).

### Voxel- and Region-of-interest-based Lesion-Symptom Mapping

Lesion overlap distributions for the entire cohort are illustrated in **Figure 1** and **eFigure 1**. The statistical results of the VLSM and ROI-LSM analyses are presented in **Figure 2** and **Table 2**.

**Figure 1.**
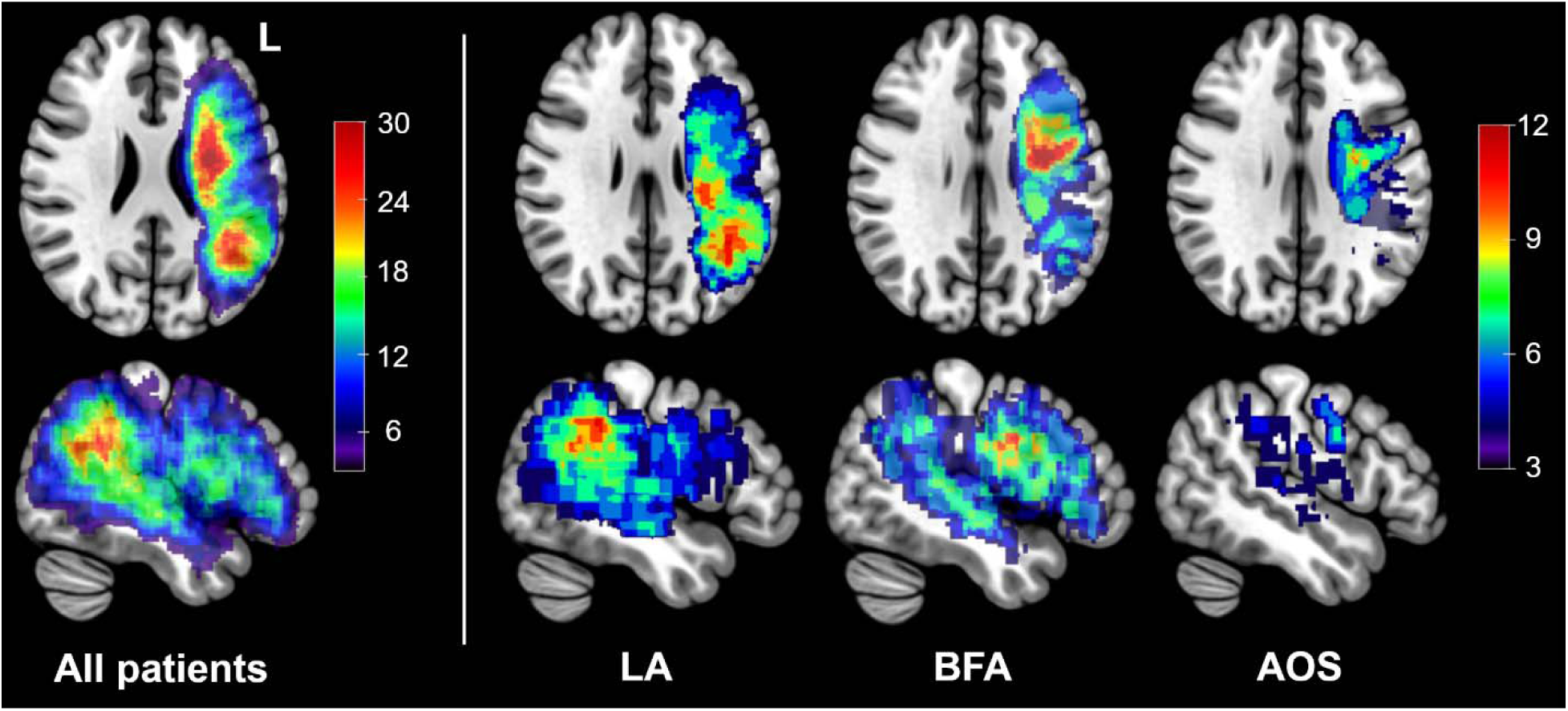
Lesion distributions of the study cohort. Color bars indicate the number of overlapping lesions for all patients and each apraxia subtype. LA: limb apraxia; BFA: buccofacial apraxia; AOS: apraxia of speech.

**Figure 2.**
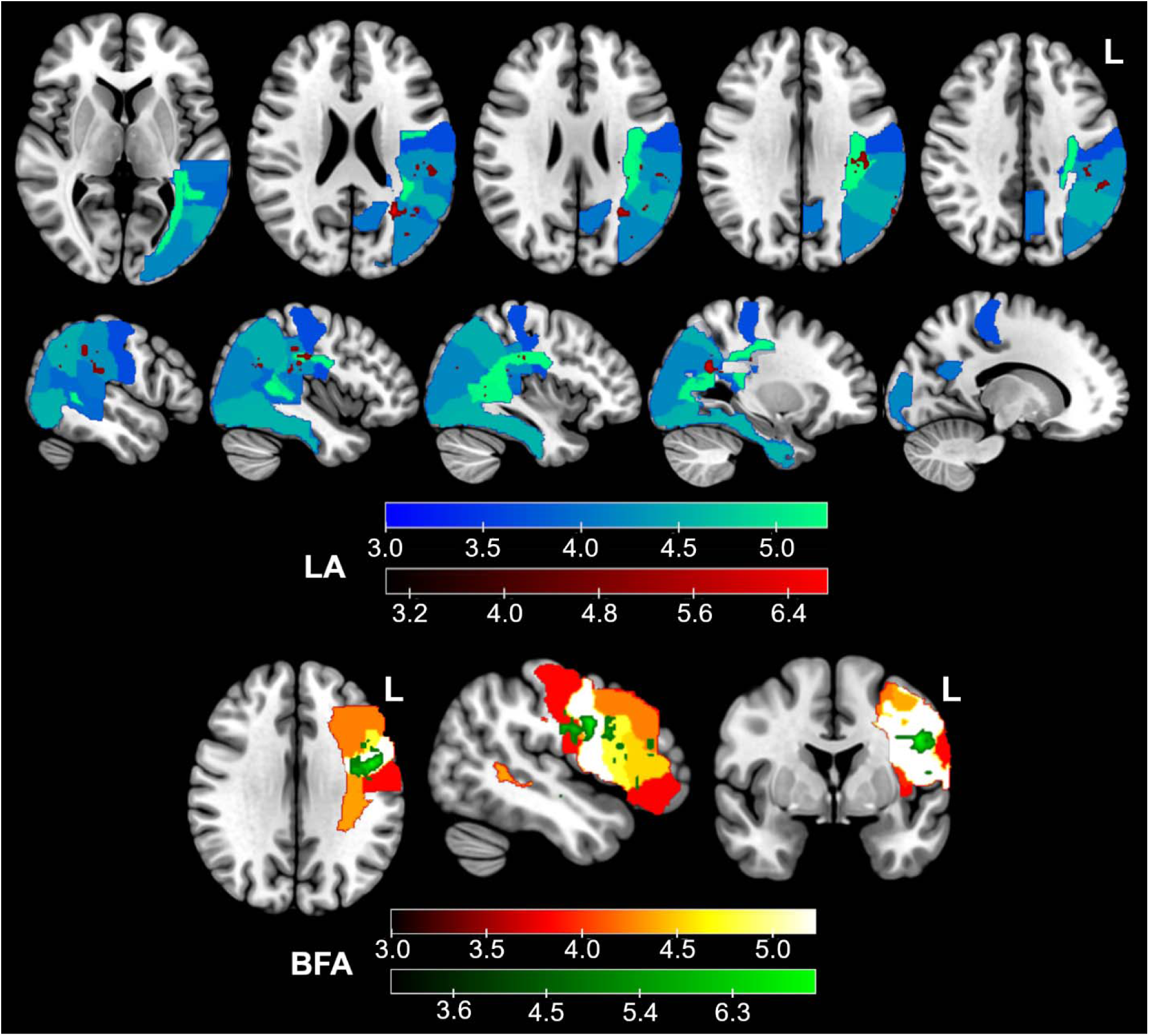
Results of voxel- and region-of-interest-based lesion-symptom mapping. The upper row displays cortical and subcortical regions significantly associated with limb apraxia (LA); VLSM results are shown in red, and ROI-LSM results in blue-green. The lower row displays regions associated with buccofacial apraxia (BFA); VLSM results are shown in green, and ROI-LSM results in red-orange-white. Color bars indicate *z*-scores. The family-wise error-corrected threshold of *p* < 0.05 corresponds to *z* > 4.9 (VLSM) and *z* > 3.6 (ROI-LSM) for LA, and *z* > 4.8 (VLSM) and *z* > 3.7 (ROI-LSM) for BFA. No suprathreshold regions were identified for apraxia of speech (*z* > 4.8 for VLSM; *z* > 3.5 for ROI-LSM).

**Table 2.** Significant regions identified by region-of-interest-based lesion-symptom mapping.

| ROIs | z-score |
| --- | --- |
| **LA** |  |
| Superior longitudinal fasciculus | 5.26 |
| Posterior thalamic radiation | 5.17 |
| Fusiform gyrus | 4.57 |
| Inferior occipital gyrus | 4.54 |
| Angular gyrus | 4.53 |
| Supramarginal gyrus | 4.28 |
| Middle occipital gyrus | 4.19 |
| Precuneus | 4.05 |
| Posterior superior temporal gyrus | 4.03 |
| Tapetum of corpus callosum | 3.96 |
| Posterior middle temporal gyrus | 3.92 |
| Postcentral gyrus | 3.66 |
| **BFA** |  |
| Precentral gyrus | 5.22 |
| Inferior frontal gyrus, pars opercularis | 4.76 |
| Inferior frontal gyrus, pars triangularis | 4.53 |
| Superior longitudinal fasciculus | 4.35 |
| Middle frontal gyrus | 4.25 |
| Meynert nucleus | 4.24 |
| Sagittal stratum | 4.20 |
| Anterior corona radiata | 4.17 |
| Insula | 3.94 |
| Postcentral gyrus | 3.84 |
| Inferior frontal gyrus, pars orbitalis | 3.81 |
| **AOS** |  |
| No suprathreshold regions |  |

Regions are listed in descending order of z-scores. The family-wise error-corrected threshold of $p < 0.05$ corresponds to $z > 3.6$ for LA, $z > 3.7$ for BFA and $z > 3.5$ for AOS.
Abbreviations: AOS, apraxia of speech; BFA, buccofacial apraxia; LA, limb apraxia; ROI, region of interest.

Regarding LA, VLSM revealed an association with a distributed pattern of damage involving both cortical structures and deep white matter in the posterior temporo-parietal region. Cortical involvement included the supramarginal, angular, and postcentral gyri. Beneath these cortical regions, significant clusters extended into the centrum semiovale, corresponding to the superior longitudinal fasciculus/arcuate fasciculus (SLF/AF), and into the deep white matter adjacent to the trigone of the lateral ventricle (forceps major and tapetum of the corpus callosum). ROI-LSM analyses corroborated these findings, confirming significant associations with the SLF/AF, posterior thalamic radiation, tapetum, and widespread posterior cortical areas including the precuneus and temporo-occipital regions.

In contrast to the distributed posterior network observed in LA, buccofacial apraxia (BFA) was associated with a more focal anterior lesion pattern centered on the ventral sensorimotor cortex. VLSM identified significant clusters localized primarily to the mid-lower portion of the left precentral gyrus, with extensions into the adjacent postcentral gyrus and the middle and inferior frontal gyri. This focal anterior topography was supported by ROI-LSM, which implicated the same sensorimotor and frontal cortical regions alongside the insula. Furthermore, ROI-LSM indicated the involvement of underlying white matter tracts, specifically the anterior corona radiata, sagittal stratum (encompassing the inferior longitudinal and fronto-occipital fasciculi), and the SLF/AF.

For apraxia of speech (AOS), neither VLSM nor ROI-LSM analyses revealed any suprathreshold voxels or regions significantly associated with the deficit at the population level after correcting for multiple comparisons.

### Connectome-based Lesion-Symptom Mapping

The results of the CLSM analyses are shown in **Figure 3**. LA was characterized by extensive structural disconnections within a posterior network. Significant disconnections involved the superior longitudinal fasciculus/arcuate fasciculus (SLF/AF), posterior thalamic radiation (PTR), and massive interhemispheric callosal fibers traversing the splenium. These tracts link widespread posterior cortical areas, including the supramarginal, angular, superior parietal, and posterior temporal cortices. Milder disconnection was also observed in the superior fronto-parietal network.

**Figure 3.**
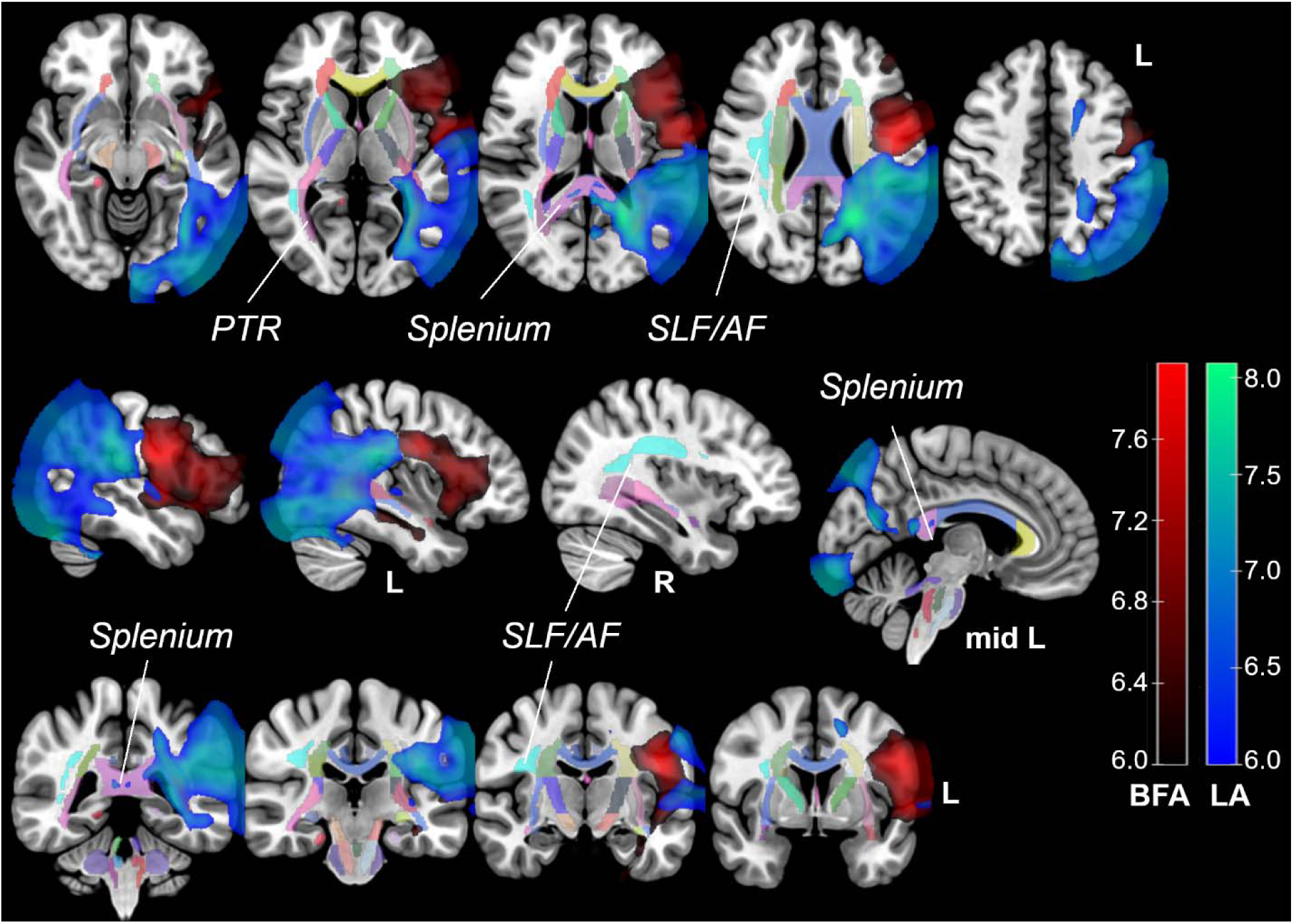
Results of connectome-based lesion-symptom mapping. Statistical disconnection maps significantly associated with limb apraxia (LA) are shown in blue-green, and those associated with buccofacial apraxia (BFA) are shown in red. The results are superimposed on the Johns Hopkins University (JHU) white matter atlas.　Color bars indicate *z*-scores. The family-wise error-corrected threshold of *p* < 0.05 corresponds to *z* > 5.9 for both LA and BFA. No suprathreshold disconnections were identified for apraxia of speech (*z* > 5.8). PTR: posterior thalamic radiation; SLF/AF: superior longitudinal fasciculus/arcuate fasciculus.

BFA was associated with focal disconnection centered on the mid-lower portion of the left precentral gyrus, extending to the inferior and middle frontal gyri and the postcentral gyrus. Unlike LA, BFA showed no significant involvement of long-range intrahemispheric or interhemispheric callosal disconnections.

For AOS, the CLSM analysis did not reveal any suprathreshold disconnections. To further explore this negative result, a CLSM analysis using a lenient statistical threshold is provided in **eFigure 2**. This exploratory analysis localized AOS to the mid-portion of the precentral gyrus. Topologically, this cluster was situated slightly dorsal to the core region of BFA.

## DISCUSSION

This study demonstrates that effector-specific apraxias are governed by distinct principles of structural network organization.LA relies on a widely distributed, bilaterally integrated temporo-parietal network, whereas BFA is associated with focal disruptions within an anterior motor-premotor network.

### The Distributed Network of Limb Apraxia

Our results indicate that LA is not merely a consequence of focal cortical dysfunction but represents an extensive structural disconnection syndrome. VLSM and ROI-LSM analyses confirmed the critical involvement of the left supramarginal gyrus, angular gyrus, and adjacent posterior cortices. Crucially, the CLSM analysis revealed that this cortical pathology is structurally linked to massive disconnections of underlying white matter tracts, including the superior longitudinal fasciculus/arcuate fasciculus (SLF/AF) and the posterior thalamic radiation.

Functionally, the left inferior parietal cortex, particularly the supramarginal gyrus, is considered to store "motor engrams" or "motor memories", the spatiotemporal kinematic patterns of learned actions^3,18^. These abstracted representations serve as internal predictive templates grounded in prior visual and proprioceptive experience^19^. Actual motor execution relies on these stored memories to guide the online predictive modulation of upper limb actions. The extensive intrahemispheric disconnections observed here likely sever the association fibers linking the posterior multimodal network, disrupting the integration of these predictive templates into ongoing motor control. Furthermore, our CLSM results highlighted significant interhemispheric disconnections traversing the splenium of the corpus callosum. Because the left parietal cortex serves as the dominant repository of motor engrams for both upper limbs^3,18^, callosal disconnection isolates the right hemisphere from these left-lateralized representations^1,20^. This may prevent the contralateral motor system from accessing the predictive templates necessary for skilled action.

This structural perspective also provides a mechanistic explanation for a historical discrepancy in behavioral neurology. While our results anchor BFA to the anterior motor-premotor network, classic studies frequently observed BFA following parietal lesions or in association with conduction aphasia^1,21^. Our data clarify this classical observation and the frequent clinical comorbidity of LA and BFA. We found that SLF/AF damage was prominent in patients exhibiting both LA and BFA but was largely spared in isolated LA cases (**eFigure 1**). This suggests that massive fronto-parietal disconnection via the SLF/AF in large middle cerebral artery strokes contributes to the simultaneous breakdown of both limb and oral praxis. The classical attribution of BFA to posterior damage likely reflected the unquantified involvement of these extensive white matter tracts, rather than a true parietal origin for oral praxis^4^.

### Localized Cortical Networks of Oral Apraxias

In contrast to the distributed posterior network of LA, BFA was associated with localized anterior lesions centered on the mid-lower portion of the left precentral gyrus, extending into the adjacent postcentral gyrus and frontal operculum. This focal topology aligns with the notion that orofacial actions rely heavily on localized intrinsic motor-proprioceptive loops^22^, supported by these adjacent motor and somatosensory cortices, rather than on extensive multisensory integration.

Regarding AOS, our primary LSM analyses did not yield suprathreshold neuroanatomical correlates. This negative finding must be interpreted within the context of our inclusive analytical approach. In our previous study using the same acute stroke cohort^9^, we demonstrated a clear anatomical dissociation between pure AOS and AOS accompanied by aphasia. While pure AOS is specifically associated with focal lesions in the mid-portion of the precentral gyrus, a localization further supported by recent studies^9,23–25^, AOS with aphasia is driven by massive, heterogeneous perisylvian lesions extending into the insula and inferior frontal gyrus^9,26,27^. When using an inclusive GLM framework that encompasses the full clinical variability of the current cohort, the spatially restricted signal of pure AOS is likely washed out by the high spatial variance of these massive concomitant strokes. Indeed, when a lenient exploratory threshold was applied (**eFigure 2**), a cluster associated with AOS emerged in the mid-portion of the precentral gyrus.

The anatomical observation that the exploratory AOS cluster is situated slightly dorsal to the core region of BFA (**eFigure 2**) is consistent with the dual organization of the laryngeal motor cortex within the left precentral gyrus^25,28,29^. Pure AOS has been consistently localized to the mid-portion of the precentral gyrus, corresponding to the dorsal laryngeal motor cortex9,23–25. In contrast, BFA localizes just below this area, centering on the mid-lower portion of the precentral gyrus and extending into the adjacent frontal operculum, which corresponds to the ventral laryngeal motor cortex^30^. Anatomically, the dorsal part of the precentral gyrus is predominantly occupied by the primary motor cortex, whereas its ventral part largely comprises the premotor cortex^31,32^. Based on this cytoarchitectonic gradient, the dorso-ventral dissociation suggests that pure AOS represents a primary motor symptom, conceptually akin to a cortical dysarthria^33^, whereas BFA represents a premotor deficit. This functional distinction aligns with direct cortical stimulation studies, where stimulation of the mid-level precentral gyrus primarily elicits positive motor responses (a primary motor phenomenon), while ventral stimulation produces negative motor responses characterized by movement inhibition^34,35^. The difference in lesion volumes observed in our cohort (**Table 1**), which were significantly larger in BFA than in AOS, further corroborates this dichotomy, as more primary cortical functions generally require smaller structural lesions to produce observable clinical deficits.

### Limitations

Several limitations should be considered. First, the use of indirect CLSM relies on a normative healthy white matter atlas rather than patient-specific tractography. While this approach avoids the tracking artifacts commonly encountered in pathological brains, it cannot account for individual premorbid anatomical variability. Furthermore, indirect disconnection estimates carry a risk of overestimation due to the assumption of complete tract disruption within the lesion mask^12^. However, it is crucial to distinguish this structural approach from functional lesion network mapping, which has recently been shown to suffer from systematic methodological biases^36^. In contrast, indirect structural disconnection measures have been demonstrated to robustly correlate with post-stroke behavioral deficits^37^, ensuring the reliability of our neuroanatomical findings.

Second, our clinical assessments had methodological constraints. Importantly, our initial clinical screening focused on patients presenting with apparent language or behavioral symptoms, introducing a potential ascertainment bias. Consequently, patients with isolated or subtle apraxias might have been underrepresented. Furthermore, because apraxias were evaluated using a binary classification (present/absent) in the acute stroke setting, the continuous severity of the deficits was not captured. This categorical approach potentially reduces statistical power. Moreover, it precluded the analysis of potential dissociations between different modes of action elicitation, such as imitation versus pantomime to verbal command, which are known to engage partially distinct neural networks^3,38,39^. Future studies employing continuous kinematic assessments and task-specific behavioral paradigms are needed to validate and extend our findings.

### Conclusions

This study provides a structural framework for understanding the distinct network architectures underlying effector-specific apraxias. We demonstrate that limb praxis relies on an extensive, bilaterally integrated temporo-parietal network, where widespread structural disconnections disrupt the distributed coordination of upper limb actions. Conversely, oral praxis depends on localized anterior cortical networks. Within the left precentral gyrus, non-verbal actions (BFA) engage a ventral premotor network, while verbal articulation (AOS) maps onto a more dorsal primary motor representation. These findings move beyond simple cortical somatotopy, highlighting how the brain uses fundamentally distinct principles of network organization, distributed versus localized, to control different classes of skilled action.

## ABBREVIATIONS

AF: arcuate fasciculus
AOS: apraxia of speech
BFA: buccofacial apraxia
CLSM: connectome-based lesion-symptom mapping
LA: limb apraxia
NIHSS: National Institutes of Health Stroke Scale
ROI: region of interest
SLF: superior longitudinal fasciculus
SLTA: Standard Language Test of Aphasia
VLSM: voxel-based lesion-symptom mapping.

## AUTHOR CONTRIBUTIONS

**Yoshiyuki Nishio:** Design or conceptualization of the study; Analysis or interpretation of the data; Drafting or revising the manuscript for intellectual content. **Ryo Itabashi:** Design or conceptualization of the study; Major role in the acquisition of data; Revising the manuscript for intellectual content. **Yuka Kataoka:** Major role in the acquisition of data. **Yukako Yazawa:** Major role in the acquisition of data. **Etsuro Mori:** Design or conceptualization of the study; Revising the manuscript for intellectual content.

## DATA AVAILABILITY

The data that support the findings of this study are available from the corresponding author, upon reasonable request. The data are not publicly available due to privacy or ethical restrictions.

## Supplementary materials

**eTable 1.** Demographic and clinical characteristics of mutually exclusive clinical subgroups.

|  | LA vs. BFA Subgroups |  |  | BFA vs. AOS Subgroups |  |  |
| --- | --- | --- | --- | --- | --- | --- |
|  | LA+/BFA-<br>(n = 12) | LA-/BFA+<br>(n = 15) | LA+/BFA+<br>(n = 5) | BFA+/AOS-<br>(n = 11) | BFA-/AOS+<br>(n = 13) | BFA+/AOS+<br>(n = 9) |
| NIHSS (Mean ± SD) | 12.7 ± 10.3 | 9.3 ± 7.1 | 16.0 ± 8.3 | 10.0 ± 6.7 | 7.7 ± 8.0 | 12.1 ± 9.2 |
| Lesion volume (cm<sup>3</sup>, Mean ± SD) | 56.4 ± 54.1 | 56.9 ± 51.5 | 159.4 ± 136.5 | 68.0 ± 44.2 | 15.6 ± 18.1* | 100.3 ± 125.7 |
| Aphasia type |  |  |  |  |  |  |
| Broca | 2 | 5 | 3 | 0 | 7 | 8 |
| Pure AOS | 0 | 1 | 0 | 0 | 6 | 1 |
| Wernicke | 7 | 4 | 2 | 6 | 0 | 0 |
| Conduction | 0 | 0 | 0 | 0 | 0 | 0 |
| Transcortical sensory | 1 | 1 | 0 | 1 | 0 | 0 |
| Anomic | 1 | 2 | 0 | 2 | 0 | 0 |
| Others | 0 | 2 | 0 | 2 | 0 | 0 |
| No aphasia or AOS | 1 | 0 | 0 | 0 | 0 | 0 |
| Comorbid apraxia |  |  |  |  |  |  |
| BFA | - | - | - | - | - | - |
| AOS | 2 | 6 | 3 | - | - | - |
| LA | - | - | - | 2 | 2 | 3 |

\* $p = 0.009$ compared to the BFA+/AOS- group.
Abbreviations: AOS, apraxia of speech; BFA, buccofacial apraxia; LA, limb apraxia;
NIHSS, National Institutes of Health Stroke Scale; (+), symptom present; (-), symptom absent.

**eFigure 1.**
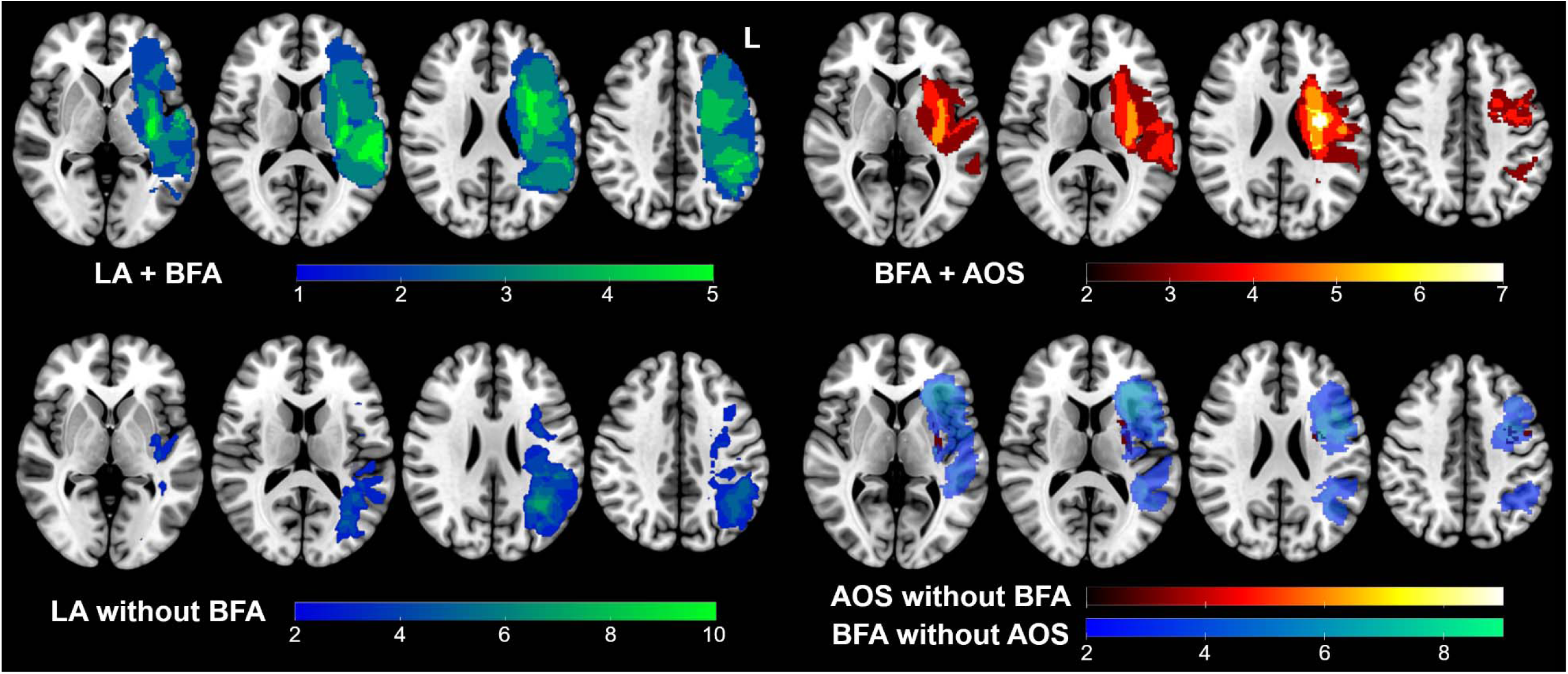
Lesion distributions of mutually exclusive clinical subgroups. Color bars indicate the number of overlapping lesions within each subgroup. LA: limb apraxia; BFA: buccofacial apraxia; AOS: apraxia of speech.

**eFigure 2.**
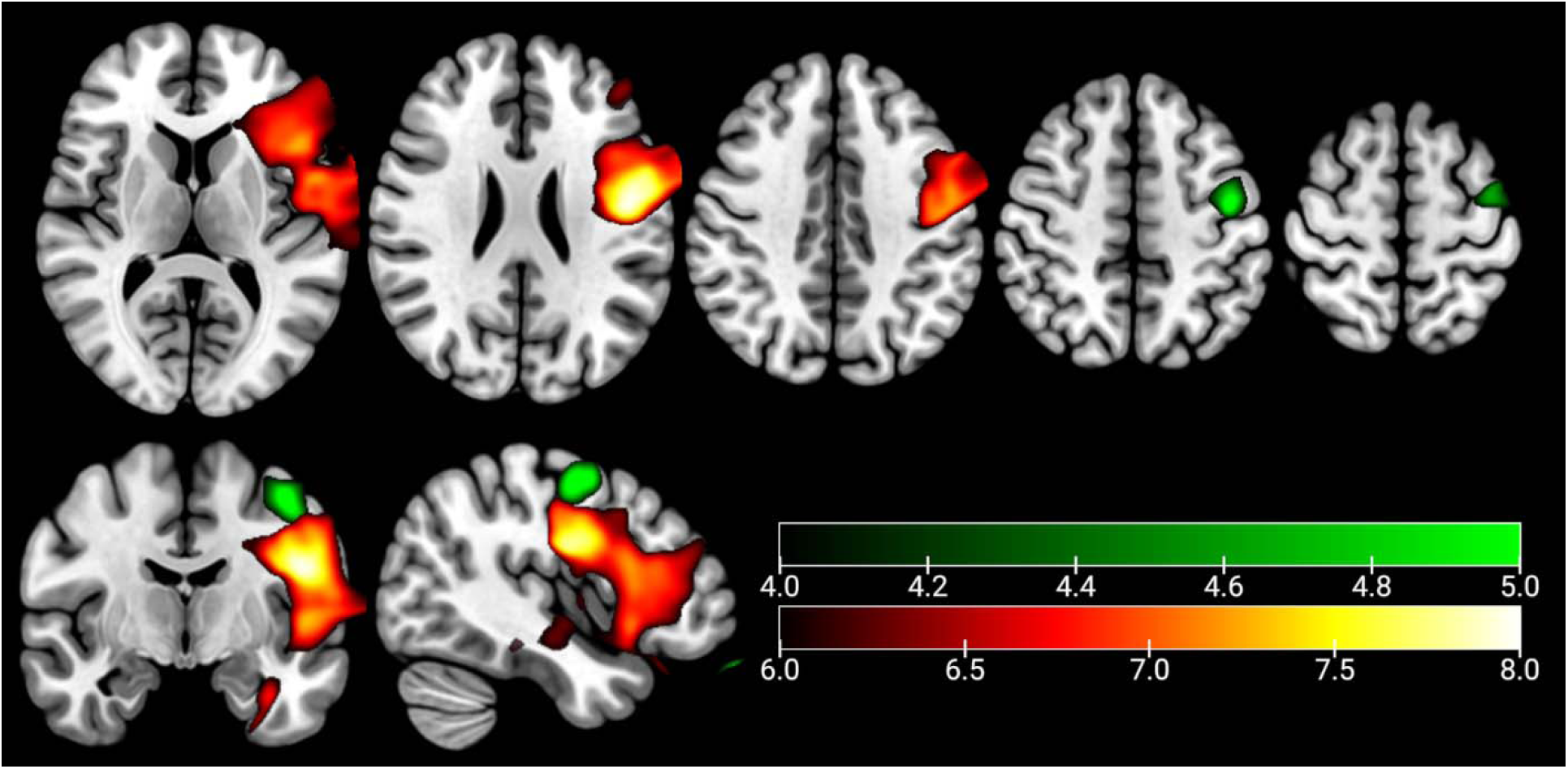
Connectome-based lesion-symptom mapping for apraxia of speech at a lenient threshold. Statistical disconnection maps associated with apraxia of speech (AOS) are shown in green using an exploratory, uncorrected threshold (*z* > 4.0). For spatial reference, the fully thresholded map for buccofacial apraxia (BFA) is overlaid in red-yellow. Color bars indicate *z*-scores.

